# Modulation of the JAK2-STAT3 pathway promotes expansion and maturation of human iPSCs-derived myogenic progenitor cells

**DOI:** 10.1101/2024.12.09.624203

**Authors:** Luca Caputo, Cedomir Stamenkovic, Matthew T. Tierney, Maria Sofia Falzarano, Rhonda Bassel-Duby, Alessandra Ferlini, Eric N. Olson, Pier Lorenzo Puri, Alessandra Sacco

## Abstract

Generation of *in vitro* induced pluripotent cells (hiPSCs)-derived skeletal muscle progenitor cells (SMPCs) holds great promise for regenerative medicine for skeletal muscle wasting diseases, as for example Duchenne Muscular Dystrophy (DMD). Multiple approaches, involving ectopic expression of key regulatory myogenic genes or small molecules cocktails, have been described by different groups to obtain SMPC towards cell-transplantation *in vivo* as a therapeutic approach to skeletal muscle diseases. However, hiPSCs-derived SMPC generated using transgene-free protocols are usually obtained in a low amount and resemble a more embryonal/fetal stage of differentiation. Here we demonstrate that modulation of the JAK2/STAT3 signaling pathway during an *in vitro* skeletal muscle differentiation protocol, increases the yield of *PAX7+* and *CD54+* SMPCs and drive them to a post- natal maturation stage, in both human ES and patient-derived iPSCs. Importantly, upon removal of the inhibition from the cultures, the obtained SMPCs are able to differentiate into multinucleated myotubes *in vitro.* These findings reveal that modulation of the JAK2/STAT3 signaling pathway is a potential therapeutic avenue to generate SMPCs *in vitro* with increase potential for cell-therapy approaches.

## Introduction

Generation of human pluripotent stem cell (hiPSCs)-derived skeletal muscle progenitor cells (SMPCs) *in vitro* holds great potential in regenerative medicine as a therapeutic approach for muscle wasting conditions, such as Duchenne Muscular Dystrophy (DMD) or sarcopenia^1,2^. Differentiation protocols that rely on ectopic expression of myogenic transcription factors, such as MYOD1/BAF60c^3^, PAX7^4^, or PAX3^5^, while they generate a homogeneous progeny of differentiated cells suitable for *in vitro* disease modeling, they possess the risk of random integration of exogenous viral/plasmid DNA, hindering their use in cell therapy^6^. On the other hand, transgene-free directed differentiation of hiPSC-derived SMPCs results in a progeny that is heterogenous and immature compared to their postnatal counterparts, hindering their utilization for *in vitro* disease modeling and/or *in vivo* treatment of muscle diseases^7–16^.

The JAK/STAT pathway has an important role in multiple embryonic and adult stem cells, regulating self-renewal, differentiation, and proliferation^17–25^. It has been previously described that the JAK2/STAT3 pathway plays a crucial role in skeletal muscle and regulates the function of several cell types, including muscle stem cells (MuSCs, also known as satellite cells) and myofibers^26^. In skeletal muscle, STAT3 mediates interleukins (IL-6, IL-10, IL-11, Oncostatin M) and other cytokines signaling (TGFβ1)^27^. Upon pathway activation, STAT3 is phosphorylated on tyrosine 705 (pSTAT3^Y705^), that induces dimerization and nuclear translocation, via the action of various kinases, with JAKs (Janus Kinases) being the main mediators of this phosphorylation event on STAT3^26^. We and others have previously demonstrated that the JAK2/STAT3 signaling axis has a critical role in regulating symmetric cell division, proliferation, differentiation, metabolic switch and autophagy in murine MuSCs, as well as in primary human myoblasts^28–32^. In our previous study we provided evidence that pSTAT3^Y705^ induces *Myod1* expression in MuSCs, promotes myogenic commitment and antagonizes self-renewal^29^. Increased IL-6 signaling and pSTAT3^Y705^ levels have been observed in aged tissues and in numerous muscle-wasting diseases, leading to muscle atrophy^30,33–37^. Transient JAK2/STAT3 inhibition *in vivo*^29,30^, or conditional MuSC specific Stat3 ablation (*Pax7^CreER/+^:Stat3^loxP/loxP^*)^29^ promotes expansion of MuSCs and muscle repair in both aged and dystrophic muscles, however the constitutive MuSC specific Stat3 ablation (*Dmd^-/-^:Stat3^loxP/loxP^:Pax7^Cre/+^*) leads to precocious differentiation and defective proliferation due to *Pax7* downregulation in MuSCs^38^, suggesting that modulation of the JAK2/STAT3 signaling pathway in both term of intensity and timing is critical to regulate MuSCs biology. Multiple JAK inhibitors (JAKi) have been developed and are currently approved for treatment of tumors and inflammatory diseases since the report from Meydan et al. of AG490 as a JAKi with antileukemic activity^39,40^. The therapeutic potential of AG490 has expanded to several pathological conditions such as Keloid disease^41^, brain hemorrhage^42^, and liver disease^43^.

To improve the generation of SMPCs *in vitro*, increasing the yield of SMPCs and promoting the maturation from embryonal stage to post-natal stage, we reasoned that modulation of the JAK2/STAT3 signaling axis, using small molecule compounds, would be beneficial due to its ability to promote PAX7+ cells’ expansion^29^. Here we show that transient treatment of myogenic cultures with the JAK2/3 inhibitor AG490 leads to a 2-fold increase of PAX7+ SMPCs across multiple pluripotent cell lines and the progression of the SMPCs to a postnatal stage that can be easily purified using one single surface marker, CD54/ICAM-1.

## Results

### Inhibition of JAK2/STAT3 pathway blocks human myoblast differentiation

To investigate the function of the JAK2/STAT3 pathway in primary healthy and dystrophic human myoblasts, we evaluated the ability of different JAK1/2 inhibitors (JAKi) to prevent Myogenin expression and myotube formation. Treatment of human primary myoblasts at 3 days of differentiation with either JAK2/JAK3 and JAK1/JAK2 inhibitors (AG490 and Ruxolitinib, respectively) or with JAK2 specific inhibitor (Gandotinib), but not with a JAK1 specific inhibitor (Filgotinib), resulted in loss of STAT3 phosphorylation - pSTAT3^Y705^ -, but only AG490 and Gandotinib treatments resulted in downregulation of Myogenin expression (**Figure 1A**). AG490 treatment completely ablated myotube formation *in vitro* of human primary myoblasts from healthy donors and Duchenne Muscular Dystrophy (DMD) patients as demonstrated by the absence of multinucleated myosin heavy chain- positive myotubes (**Figure 1B**) and the decrease of the differentiation index, calculated as the percentage of nuclei in myosin heavy chain-positive cells, in a dose dependent manner (**Figure 1C**). Treatment of healthy and DMD primary myoblasts with 10 µM AG490 resulted in a noticeable decrease of pSTAT3^Y705^ signal (**Figure 1D**), and promoted myoblasts proliferation (**Figure 1E-F**). We next decided to test whether other JAKi had similar effects on myotubes formation in DMD-derived myoblasts and observed a block of myogenic differentiation when the specific JAK2i Gandotinib was added to myogenic cells (**Supp Figure 1 A-B**). However, we observed a significant increase in apoptotic cells, as shown by the percentage in TUNEL-positive cells in Gandotinib-treated samples (**Supp Figure 1C-D**). Thus, AG490 was selected as JAK2/STAT3 inhibitor for the remaining of the project. All together, these results confirmed the importance of JAK2/STAT3 signaling pathway in human myogenesis and reinforced the idea of a potential therapeutic approach to expand a population of muscle progenitor cells *in vitro*.

**Figure 1:**
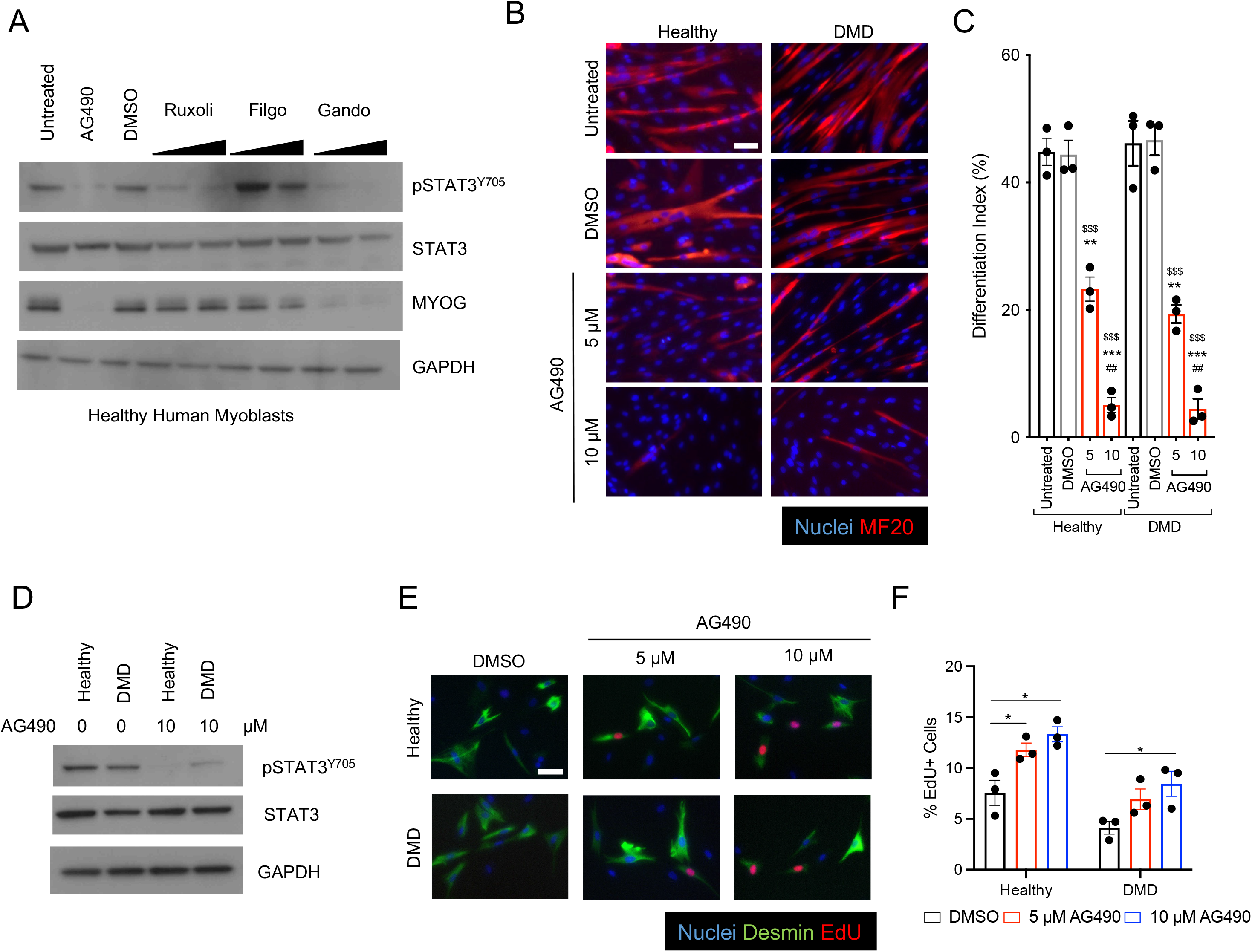
AG490 mediated inhibition of JAK/STAT3 signaling blocks differentiation of human myoblasts A. Western blot analysis of whole cell lysate of human healthy myoblasts treated with various JAK inhibitors using a-pSTAT3, a-total STAT3, anti-Myogenin and anti-GAPDH antibodies. B. Representative immunofluorescence images of myotubes either from healthy or Duchenne patient-derived myoblasts treated with AG490 or DMSO as vehicle control for MF-20 (red), nuclei are counterstained with Hoechst (blue). Scale bar 100 µm. C. Differentiation index of myotubes for the conditions shown in (B) n=3 biological replicates. D. Western blot analysis of whole cell lysate of human healthy and Duchenne patient-derived myoblasts treated with 10 µM AG490 using a-pSTAT3, a-total STAT3, and anti-GAPDH antibodies. E. Representative immunofluorescence images of myoblasts stained with Desmin (green), EdU (red) nuclei are counterstained with Hoechst (blue). Scale bar 20 µm. F. Quantification of the percentage of proliferating myoblasts treated with DMSO, 5 µM or 10 µM AG490 through EdU staining n=3 biological replicates. ns. p > 0.05 ; *. p≤ 0.05 ; **. p≤ 0.01 ; ***. p≤ 0.001 unpaired Student’s t test. * over untreated control; # over lower concentration of drug; $ over DMSO – vehicle control in C.

### AG490 treatment of stem cells-derived human myogenic cultures leads to expansion of PAX7- positive progenitor cells

To test whether JAK2/STAT3 inhibition can lead to expansion of patient-derived muscle progenitor cells *in vitro*, we decided to focus on myogenic differentiation protocols that rely on developmental signals, chemical defined, xeno-free media without the ectopic expression of myogenic transcription factors. We initially set out to evaluate two myogenic protocols – Shelton et al (Protocol 1)^9^ and Chal et al (Protocol 2)^10^ – based on the survival of cultures in our hands and efficiency of mesoderm generation we decided to test JAK2/STAT3 inhibition in Protocol 2. To understand the time window when the STAT3 pathway is active during the *in vitro* hESCs (H9) myogenic differentiation protocol, we monitored the expression of *SOCS3*, a direct transcriptional target of pSTAT3^29,30^. We observed upregulation of *SOCS3* transcription at day 12 of the differentiation protocol (**Figure 2A**), coinciding with appearance of PAX3+ paraxial mesodermal progenitors (**Figure 2B**), compatible with the essential role of STAT3 for mesoderm development during embryogenesis^44^, and sustained *SOCS3* expression up to days 25-30, when myoblasts, muscle progenitors and myocytes are present in the culture condition^11^. We treated muscle cultures with different concentrations of AG490 (2.5 and 5 µM) starting from day 12 to day 30 (**Figure 2B**) and observed continuous and dose-dependent inhibition of *SOCS3* mRNA levels (**Figure 2C**). Importantly, we observed upregulation of *PAX7* mRNA levels at day 30 only when AG490 treatment was started at early stages of the myogenic process (day 12 to day 30), but not when the treatment was started at later stages, when the peak of STAT3 activity already declined (day 15 to day 30) (**Figure 2D**). We confirmed the upregulation of PAX7 by immunofluorescence (IF) analysis of day 30 cultures (**Figure 2E**). Quantification of the PAX7 and MYOG intensity staining revealed increased intensity of both PAX7 and MYOG staining in AG490- treated cultures (**Figure 2F-G**), and a significant increase (∼2 fold) of PAX7+ nuclei in AG490-treated cultures (0 µM 10.85%±0.50%; 2.5 µM 21.94%±2.95%; 5 µM 19.96%±0.70%) (**Figure 2H**). All together, these results suggest that AG490 treatment leads to expansion of *in vitro* derived Pax7+ muscle progenitor cells.

**Figure 2:**
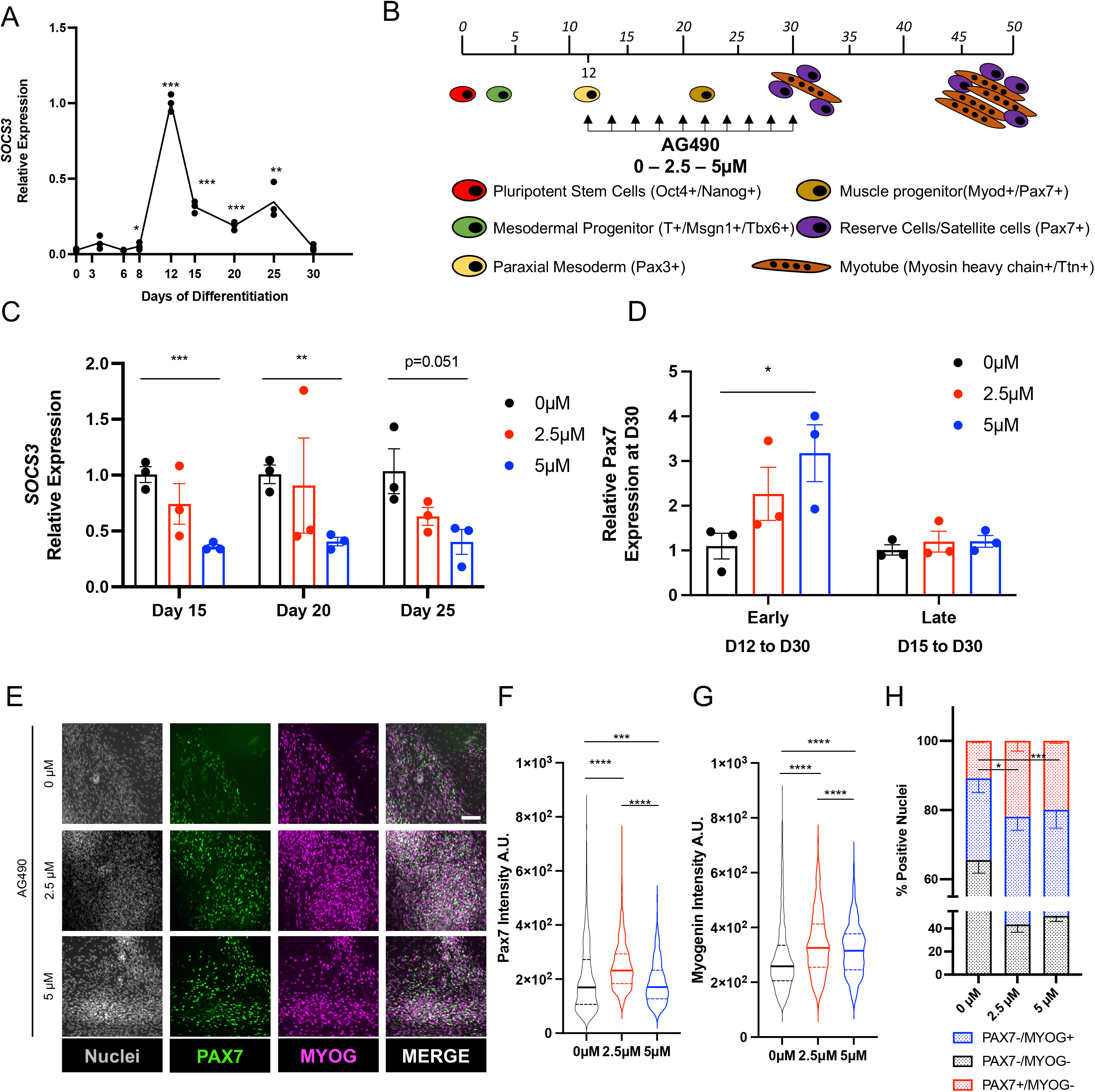
AG490 treatment leads to expansion of PAX7+ cells *in vitro* A. qPCR analysis of SOCS3 expression during myogenic differentiation. B. Schematic representation of the myogenic differentiation protocol, AG490 treatment is between D12 and D30 of differentiation, with refreshment of the media every 48hours. C. qPCR analysis of SOCS3 expression at different days of AG490 treatment. D. qPCR analysis of PAX7 expression at D30 of the differentiation protocol, with AG490 treatment started early (at D12) or late (D15). E. Representative immunofluorescence images of MYOG (magenta) and PAX7 (green) at D30 of differentiation, nuclei are counterstained with Hoechst (gray). Scale bar 50 µm. F. Violin plot or the quantification of the nuclear mean fluorescence intensity (arbitrary unit) (median ± second and third quantile) of the PAX7 IF signal staining. G. Violin plot or the quantification of the nuclear mean fluorescence intensity (arbitrary unit) (median ± second and third quantile) of the MYOG IF signal staining H. percentage of PAX7+ or MYOG+ nuclei at D30 of differentiation. N=3 biological replicates ns. p > 0.05 ; *. p≤ 0.05 ; **. p≤ 0.01 ; ***. p≤ 0.001 unpaired Student’s t test for A - C - H, One-Way ANOVA for F – G.

Next, to investigate whether myogenic cultures treated with AG490 retained the differentiation potential, we exchanged the media to differentiation media and lifted the inhibitor treatment. All myogenic cultures were able to generate myosin heavy chain-positive myotubes and PAX7+ myogenic progenitor cells at day 50 of the differentiation protocol (**Figure 3A**). AG490-treated cultures showed an upregulation of myogenic markers of both progenitors (*PAX7*) and more differentiated myocyte-like cells (*MYOD1, MYOG*) both at mRNA and a protein level (**Figure 3A-B**). Interestingly, we could observe the appearance of a higher PAX7 molecular weight band, compatible with a post-translational modification pattern for PAX7, in cultures treated with 2.5 µM AG490 compared to the control cultures, or cultures treated with 5 µM AG490 (**Figure 3C** red asterisk). This finding is of particular interest considering that hyperacetylation or phosphorylation of PAX7, modifications compatible with the migration shift observed in the western blot analysis, have been linked to an increased self-renewal capability of MuSCs^45,46^.

**Figure 3:**
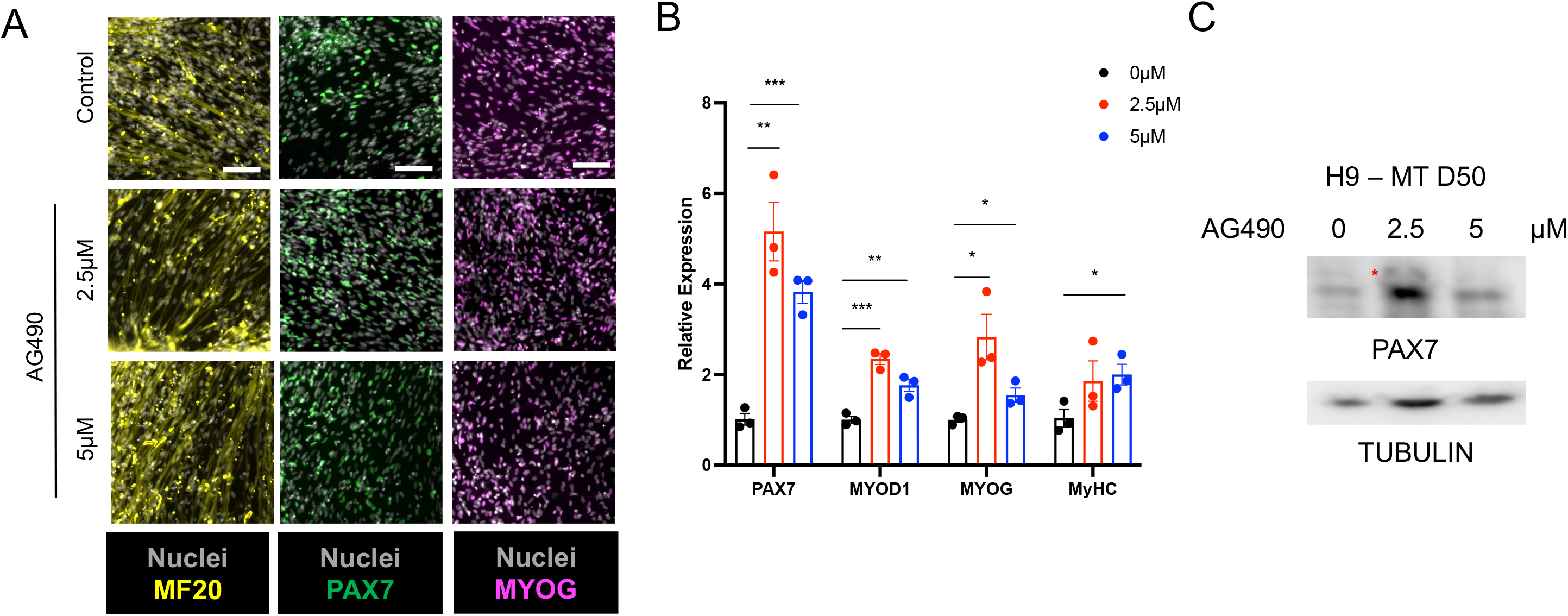
Increased myogenic differentiation upon AG490 removal A. Representative images of MF-20 (yellow), PAX7 (green) and MYOG (magenta) at D50 of differentiation, nuclei are counterstained with Hoechst (gray). Scale bar 100 µm for MF-20, 50 µm for PAX7 and MYOG. B. qPCR analysis of myogenic factors (PAX7, MYOD1, MYOG, and MyHC) expression at D50 of differentiation. C. Western blot analysis of whole cell lysate of H9-derived myogenic cultures treated with AG490 using a-PAX7, and anti-Tubulin antibodies N=3 biological replicates. ns. p > 0.05 ; *. p≤ 0.05 ; **. p≤ 0.01 ; ***. p≤ 0.001 unpaired Student’s t test.

### Generation of Pax7 positive myogenic progenitors from patient-derived hiPSCs

To evaluate the ability of AG490 to promote PAX7+ cells expansion in different patient-derived hiPSCs lines, we generated hiPSCs from a patient carrying a deletion of exon 52 of the *dystrophin* gene and from a healthy donor (**Supp Figure 2A**) using the Sendai virus system to transiently express the Yamanaka reprogramming factors^47,48^. We confirmed in two separate clones per line that expression of the pluripotency marker OCT4 was comparable to an established human embryonic cells line, H9 cell (**Supp Figure 2B**) and that when differentiated into embryoid bodies the obtained hiPSCs were able to generate all the three germ layers as detected by the expression of marker genes for each germ layer (*OTX2* – ectoderm; *SOX17* – endoderm; *MSGN* – mesoderm) (**Supp Figure 2C**). We first validated that both healthy and DMDΔ52 derived hiPSCs were able to generate PAX7+ myogenic progenitor cells with a similar efficiency (**Supp Figure 3A**). Given that 2.5 µM AG490 showed the most efficient results in H9-derived myogenic cultures, we selected this concentration in all of the subsequent experiments. AG490 treatment led to a decreased expression of *SOCS3* at day 15 of the differentiation protocol for both healthy and DMD derived hiPSCs (**Figure 4A**). Inhibition of the JAK2/STAT3 pathway in the hiPSCs-derived myogenic cultures did not completely block the progression and differentiation of the progenitor cells towards MYOG+ myocytes as we could detect generation of both PAX7+ and MYOG+ populations at day 30 of culture (**Figure 4B** and **Supp Figure 3B**). Similarly to what we observed in the H9 hESC line, we observed a ∼2 fold increase of PAX7+ nuclei in hiPSCs control, but not in DMDΔ52 (Healthy 0 µM 8.67%±1.04% vs 2.5 µM 17.97%±1.12%; DMDΔ52 0 µM 15.55%±0,92% vs 2.5 µM 18.64%±3.32%) (**Figure 4C**), in concordance with previously described roles for dystrophin in mediating signaling in muscle progenitor cells^49^. PAX7 upregulation in AG490-treated myogenic cultures was confirmed using immunofluorescence (IF) analysis at day 30 of differentiation (**Figure 4D**). Furthermore, AG490 treatment of hiPSCs-derived myogenic cultures do not completely block the progression and differentiation of the progenitor cells as we could detect increase expression of *PAX7*, *MYOG*, and *MYOD1* (**Figure 4E**). Importantly, the obtained PAX7+ SMPCs were able to differentiate into myosin heavy chain+ myotubes when switched to differentiation media expressing myogenic markers (**Figure 4F-G**). These results were validated in an independent clone of hiPSCs control and DMDΔ52 (**Supp Figure 3C-D**). Finally, since different hiPSCs lines can respond to direct differentiation protocols with different efficiency and different DMD mutations can have specific response to pharmaceutical treatments, we tested AG490 treatment on hiPSCs lines carrying a deletion of DMD exon 8 and 9 (DMDΔ8/9), leading to a premature stop codon and their isogenic control line. As observed in H9 and the previously used hiPSCs lines, control line (AMD_Healthy), and the derived line carrying DMDΔ8/9 mutation (AMD_iDMDΔ8/9)^50^ responded to AG490 treatment by increasing expression of PAX7 (**Supp Figure 4**).

**Figure 4:**
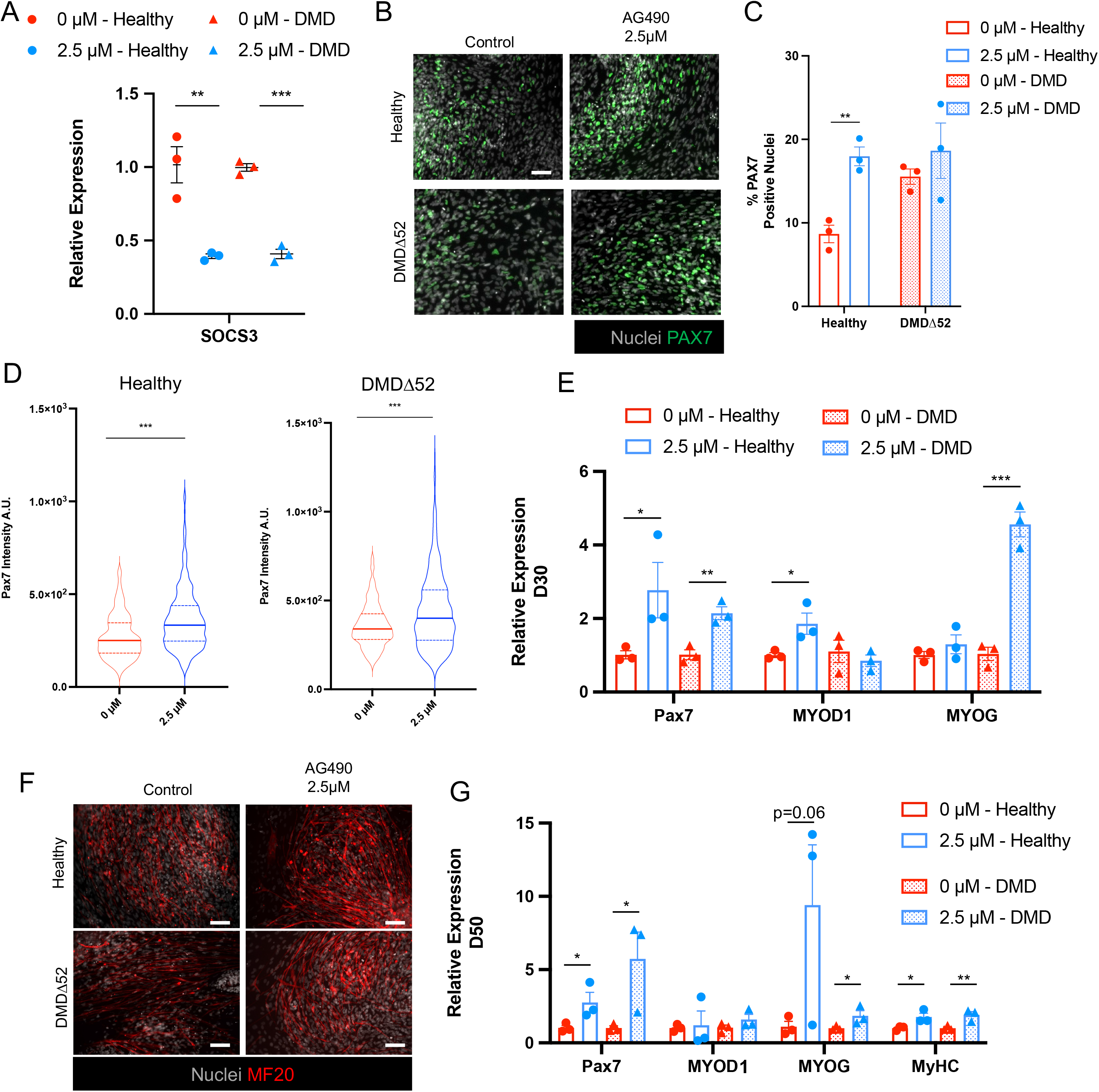
AG490 treatment leads to increased myogenesis in both healthy and DMD myogenic cultures A. qPCR analysis of SOCS3 expression at D30 of the myogenic differentiation protocol in healthy (circles) and DMDΔ52 (triangles) myogenic cultures. B. Representative immunofluorescence images of PAX7 (green) at D30 of differentiation, nuclei are counterstained with Hoechst (gray). Scale bar 50 µm. C. Percentage of PAX7+ nuclei at D30 of differentiation D. Violin plot or the quantification of the nuclear mean fluorescence intensity (arbitrary unit) (median ± second and third quantile) of the PAX7 IF signal staining in healthy (left panel) and DMDΔ52 (right panel) myogenic cultures. E. qPCR analysis of *PAX7, MYOD1* and *MYOG* expression at D30 of the differentiation protocol F. Representative immunofluorescence images of MF-20 (red) at D50 of differentiation protocol, nuclei are counterstained with Hoechst (gray). Scale bar 50 µm. G. qPCR analysis of myogenic factors (*PAX7, MYOD1, MYOG*, and *MyHC*) expression at D50 of differentiation. N=3 biological replicates. ns. p > 0.05 ; *. p≤ 0.05 ; **. p≤ 0.01 ; ***. p≤ 0.001 unpaired Student’s t test for A - C - E - G, One- Way ANOVA for D.

### JAK2/STAT3 inhibition leads to the generation of postnatal SMPCs

In order to better characterize the population of SMPCs obtained upon AG490 treatment, we decided to employ a one-step purification protocol of PAX7+ cells that relies on the expression of the surface marker CD54/ICAM-1^51^. Using a CD54 antibody in conjunction with magnetic-activated cell sorting (MACS) at day 30 of the direct differentiation protocol, we could observe a ∼2 fold increase of CD54+ cells isolated from myogenic cultures treated with 2.5 µM AG490 compared to untreated, increase that was absent in myogenic cultures treated with 5 µM AG490 (0 µM 7.50%±1.41%; 2.5 µM 16.23%±0.62%; 5 µM 6.00%±0.44%) (**Figure 5A**). As expected, isolated CD54+ cells were enriched for *CD54* mRNA expression, but only 2.5 µM treated cells showed significantly enrichment for *PAX7* (**Figure 5B**). CD54+ SMPCs from 2.5 µM-treated myogenic cultures showed enrichment for *MYOD1* and *MYOG* (**Figure 5C**), suggesting a poised activation signaling. A key limitation of in vitro generation of SMPCs is that, independently from the protocol used, the obtained SMPCs resemble at a transcriptional and fusogenic levels embryonal/fetal SMPCs rather than a postnatal stage^8,52–54^. Intriguingly, analysis of publicly available datasets^8^ revealed that CD54/ICAM-1 is specifically expressed by late stages SMPCs, a stage comprised by postnatal MuSCs (**Figure 5D**, **Supp Figure 5A-B**). We therefore decided to investigate whether AG490 treatment could induce a further maturation *in vitro* of SMPCs that resemble postnatal MuSCs at the transcriptional level. We decided to test the expression of specific markers from the previously identified developmental stages ^8^ (**Figure 5D**, **Supp Figure 5A-B**). CD54+ cells isolated from 2.5 µM treated myogenic cultures showed enrichment for markers linked to Stage4 (*NFIB*, *NCOA1,* and *PLAGL1*), while CD54+ cells isolated from cultures treated with 5 µM AG490 showed enrichment for markers linked to Stage1/2 (*PAX3, ID2,* and *SOX4*) (**Figure 5E** and **Supp Figure 5**), suggesting that regulation of JAK/STAT3 signaling is critical for expansion and maturation of SMPCs.

**Figure 5:**
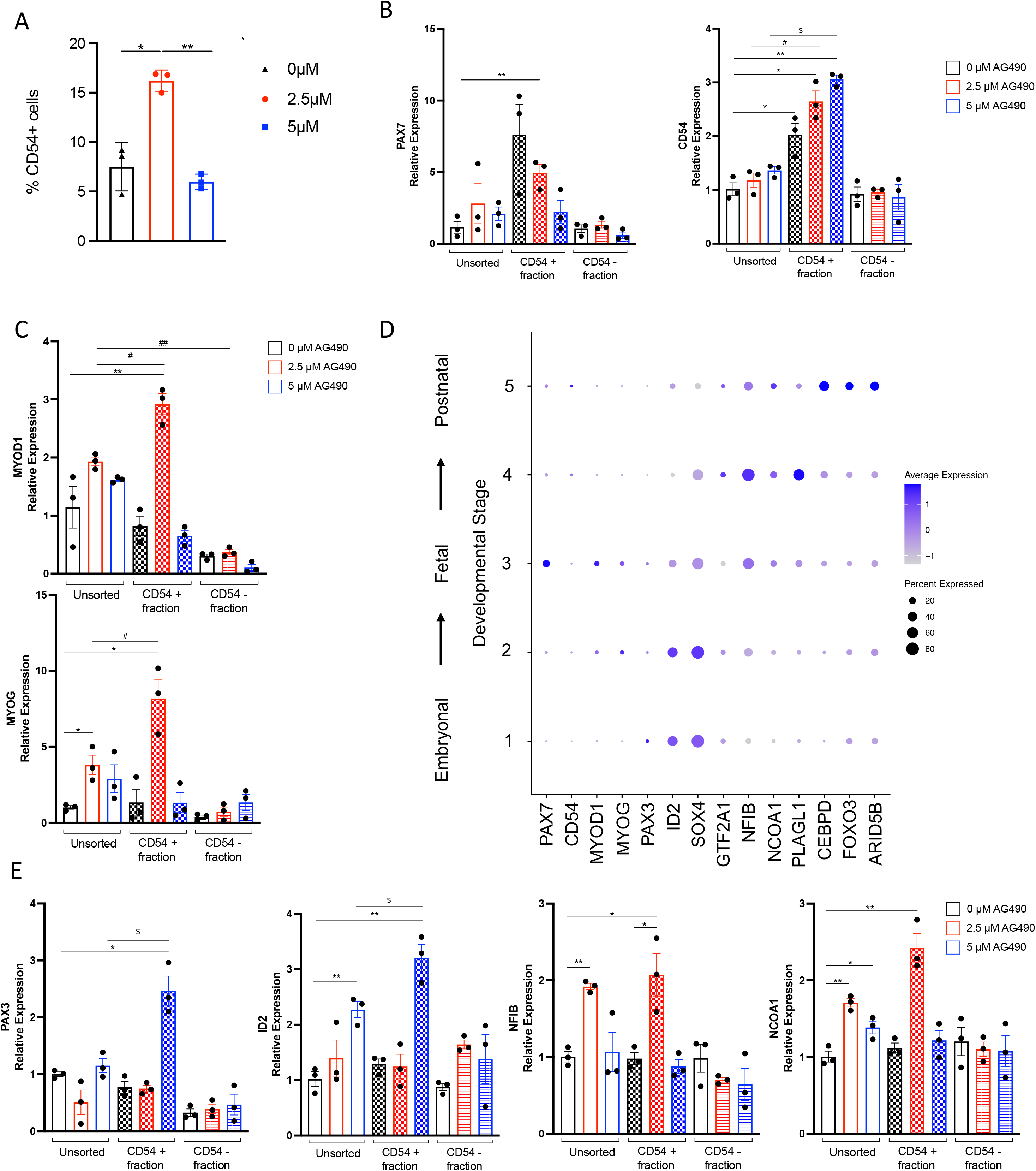
AG490 treatment leads to maturation of *in vitro* derived SMPC A. Percentage of CD54+ cells obtained from H9-derived myogenic cultures at D30 of differentiation. B. qPCR analysis of *PAX7* and *CD54/ICAM1* expression in CD54 MACS-purified populations. C. qPCR analysis of *MYOD1* and *MYOG* expression in CD54 MACS-purified populations. D. Dot plot analysis for selected transcription factors differentially expressed in embryonal (STAGE1/2) vs postnatal (STAGE 5) stages. E. qPCR analysis of *PAX3, ID2* (STAGE1/2) and *NFIB, NCOA1* (STAGE 4) expression in CD54 MACS-purified populations. N=3 biological replicates. ns. p > 0.05 ; *. p≤ 0.05 ; **. p≤ 0.01 ; ***. p≤ 0.001 unpaired Student’s t test.

## Discussion

Since the seminal works of Yamanaka group^48^ and Jaenisch group^55^, generation of hiPSCs-derived progenitor cells to be utilized in cell-replacement therapies holds great promises in regenerative medicine. We and others have developed protocols to allow the generation of myogenic cultures either by overexpression of specific transcription factors^3–5,51,56^, or by mimicking developmental cues during the process of myogenesis *in vivo*^7,9–16,57^ (reviewed in^58^). However, difficulties in derivation of SMPCs *in vitro* that are able to maintain their self-renewal capabilities, acquire a postnatal maturation stage and with a high yield have hindered the potential of these cells to be utilized as *in vitro* model of muscle diseases and muscle wasting conditions such as muscular dystrophies, sarcopenia, and cachexia. These issues also hamper the use of *in vitro* generated SMPCs for *in vivo* treatment of muscular dystrophies. Here we show that transient treatment of myogenic cultures with a small molecule inhibitor of the JAK/STAT pathway, AG490, is able to increase the yield of PAX7+ SMPCs in vitro by ∼2 fold in both hESCs and hiPSCs, enhance expression of PAX7 at both mRNA and protein levels in SMPCs, and further allow their maturation to a developmental stage that closely resembles postnatal MuSCs.

MuSCs are anatomically defined as mononucleated cells localized between the sarcolemma of the myofiber and the basal lamina^59^, molecularly marked by the expression of the transcription factor PAX7^60^, and possessing self-renewal capability *in vivo*^61^. Early transplantation experiments have revealed that MuSCs are highly heterogenous *in vivo,* more recently confirmed by scRNAseq analysis^61–65^. MuSCs with higher Pax7 expression have been linked to increased self-renewal capabilities, increased propensity for symmetric cell division, lower metabolic state, and ultimately more stemness^61,62^. Upon AG490 treatment we observed an increased expression of PAX7 at the protein level at day 30 and day 50 of differentiation when cells were treated with lower dose of JAK/STAT inhibitor, but less pronounced at higher doses. It is worth noting that while inhibition of JAK2/STAT3 pathway stimulates proliferation of MuSCs and prevents their further differentiation^29,30^, inhibition of JAK1/STAT1/STAT3 pathway induces precocious differentiation of myoblasts^66^. Therefore, it is possible to speculate that this apparently discordant phenotype observed at lower (2.5 µM) versus higher (5 µM) doses of AG490, a JAK2/JAK3 inhibitor, which at higher doses inhibits auto-phosphorylation and therefore activation of JAK1^67,68^, is attributable to the inhibition of two separate signaling pathways. Using both genetic and molecular tools, Rudnicki and colleagues demonstrated that acetylation of PAX7, tightly regulated by Myst1/KAT8 and Sirt2/SIRT2, respectively the lysine acetyltransferase and deacetylase responsible for controlling the levels of PAX7 acetylation, is necessary for MuSCs symmetric division, increased transcriptional activity of PAX7, interaction with the niche *in vivo* and higher regenerative potential of MuSCs^45^. Intriguingly, we observed a higher molecular weight band in the Western blot that is compatible with a shift caused by PAX7 acetylation. Both enzymes involved in the regulation of the acetylation status of PAX7 are bound and transcriptionally regulated by STAT3 in various cell types^69–71^. It is tempting to speculate that a conserved binding and regulation of Myst1/KAT8 and Sirt2/SIRT2 could be present also in myogenic cells. Modulation of JAK/STAT3 pathway would then be able to control the acetylation state of PAX7 and regulate activation or expansion of MuSCs in both murine animal model and in humans. Comparative scRNA-seq analysis of *in vitro* hiPSCs-generated myogenic cells with human limb skeletal muscle cells at embryonic (week 5-8), fetal (week 9-18), juvenile (year 7-11) and adult (year 34-42) stages demonstrated that *in vitro* generated myogenic cells are at an embryonic/fetal transitional stage even after several weeks in culture^8^. Multiple groups have conducted studies to improve maturation of these cultures. Inhibition of the TGFβ1 signaling pathway, alone or in combination with a NOTCH inhibitor (DAPT), corticosteroids (Dexamethasone), or activator of the adenylate cyclase (Forskolin), have been shown to promote maturation of myotubes *in vitro* with a switch from MYH3/MYH8 expression to MYH1/MYH2 and better myofibrillar organization^7,12,72–74^. However, DAPT treatment led to the differentiation of PAX7+ cells present in the myogenic culture, therefore resulting in the depletion of progenitor cells^12^. Until now, hiPSCs-derived SMPCs have been effectively matured to a more postnatal stage only upon engraftment *in vivo* in NGS-mice^75,76^. The maturation observed upon engraftment *in vivo* is potentially beneficial and demonstrate that microenvironment signals, including IL-6 and JAK2/STAT3 regulation, are necessary for maturation of MuSCs. It is, however, imperative the need to identify the signaling pathways regulating this process for the use in the clinic of *in vitro* generated SMPCs. Here, we show that upon modulation of JAK/STAT signaling pathway, in controlled *in vitro* conditions, CD54+ SMPCs present a transcriptional signature that more closely resemble the one of postnatal MuSCs (Stage 4 of the developmental trajectory). Future studies are warranted to fully understand the molecular mechanisms and the window of treatment both in term of duration and concentration of AG490, possibly in an iPS cell line-dependent manner.

In conclusion, we have demonstrated that by modulating the JAK2/STAT3 signaling pathway *in vitro* we can double the yield of PAX7+ SMPCs in both human ES and human patient-derived iPSCs. Our expression analysis also suggests that inhibition of JAK2/STAT3 signaling pathway allows PAX7+ SMPCs to mature to a postnatal stage. Accordingly, we postulate that SMPCs generated in this study could be a better model to study MuSCs behavior and potentially have an improved engrafting capability *in vivo*.

### Experimental procedures Human Cell Lines Isolation

Healthy and DMD primary myoblasts were previously described^77^. Human muscle biopsies from healthy and DMD patients were obtained from the lower extremity muscles during surgical procedures as part of the patient’s clinical care plan at Rady’s Children’s Hospital, San Diego. Written informed consent from the parent or guardian was obtained for all subjects. The protocol was approved by the University of California, San Diego Human Research Protectants Program and Institutional Review Board in accordance with the requirements of the Code of Federal Regulations on the Protection of Human Subjects. Three healthy and three DMD-affected biopsies were collected for this study.

Human and DMD skin fibroblasts were obtained at University of Ferrara. One healthy and one DMD-affected skin biopsy specimens were collected for this study. Written informed consent from the parent or guardian was obtained for all subjects. The protocol was approved by UNIFE Ethical Committee approval (Approval number 841/2020/Sper/AOUFe). Healthy iPSCs and DMDΔ52 iPSCs were generated at the Stem Cell Core at the SALK Institute from skin fibroblasts.

Anonymous male donor (AMD) iPSCs (Healthy, iDMDΔ8/9) were generated at University of Texas Southwestern Medical Center^50^.

### Cell lines maintenance and culture

Primary myoblasts were grown on collagen (BD Biosciences Cat #354254) coated dishes and maintained in growth media (GM - DMEM, 20% FBS, GlutaMAX, 10 ng/ml insulin, 24 ng/ml basic fibroblast growth factor (bFGF), 10 ng/ml epidermal growth factor (EGF) and penicillin and streptomycin). Differentiation was induced when myoblasts reached 90% confluency and media switch to differentiation media (DM – DMEM, 5% horse serum and 100 ng/ml insulin).

hES (H9 - WiCell WA09) and hiPSCs were maintained in mTESR+ media (Stem Cell Technologies Cat #100-0276) on Basement Membrane Extract (BME, Cultrex Reduced Growth Factor Basement Membrane Extract, R&D Systems Cat #3433-010-01) coated dishes in incubator at 37 °C, 5% CO_2_.

### Small molecule treatment of human primary myoblasts

AG490 (Tocris Cat #0414) was dissolved in DMSO at a concentration of 20mM. Ruxolitinib (Selleck Chemical Cat #508470) was dissolved in DMSO at a concentration of 20mM. Filgotinib (Selleck Chemical Cat #S7605) was dissolved in DMSO at a concentration of 20mM. Gandotinib (LY2784544) (Selleck Chemical Cat #S2179-5MG) was dissolved in DMSO at a concentration at a concentration of 20mM.

Healthy and DMD primary myoblasts were plated on collagen coated dishes and treated with JAKi in GM conditions for 48 hours before analysis. For proliferation analysis, myoblasts were incubated with EdU (20μM) for 4 hours at 37°C prior to fixation with 1.5% PFA for 15’ at room temperature. For fusion index analysis, myoblasts were pre-treated in GM condition with JAKi at various concentration for 48h, before switching to DM ± JAKi at specific concentration for 48h prior to fixation with 1.5% PFA for 15’ at room temperature.

### Directed differentiation protocol

hES and hiPSC lines were subjected to direct differentiation as described in Chal et al^11^. AG490 (Tocris Cat #0414) was dissolved in DMSO at a concentration of 20mM and added to differentiation media starting from D12, media was refreshed every 2 days. AG490 was maintained in media until D30 of differentiation, when the media was exchanged to N2-containing media to induce myotube differentiation.

### Gene expression analysis

Total RNA was isolated using Quick-RNA^TM^ Microprep Kit (Zymo Research Cat #R1050) according to manufacturer’s recommendation. 0.5-1 µg of total RNA was retrotranscribed using High Capacity cDNA Reverse Transcription Kit (Applied Biosystems Cat #4368813) and real-time PCR was performed using the Power SYBR Green PCR Master Mix (Applied Biosystems Cat #4367659) on Applied Biosystems 7900HT Fast Real-Time PCR. Relative gene expression was calculated using the 2^-ΔΔCT^ method and normalized to β-actin expression. Primer pairs are shown in the table below.

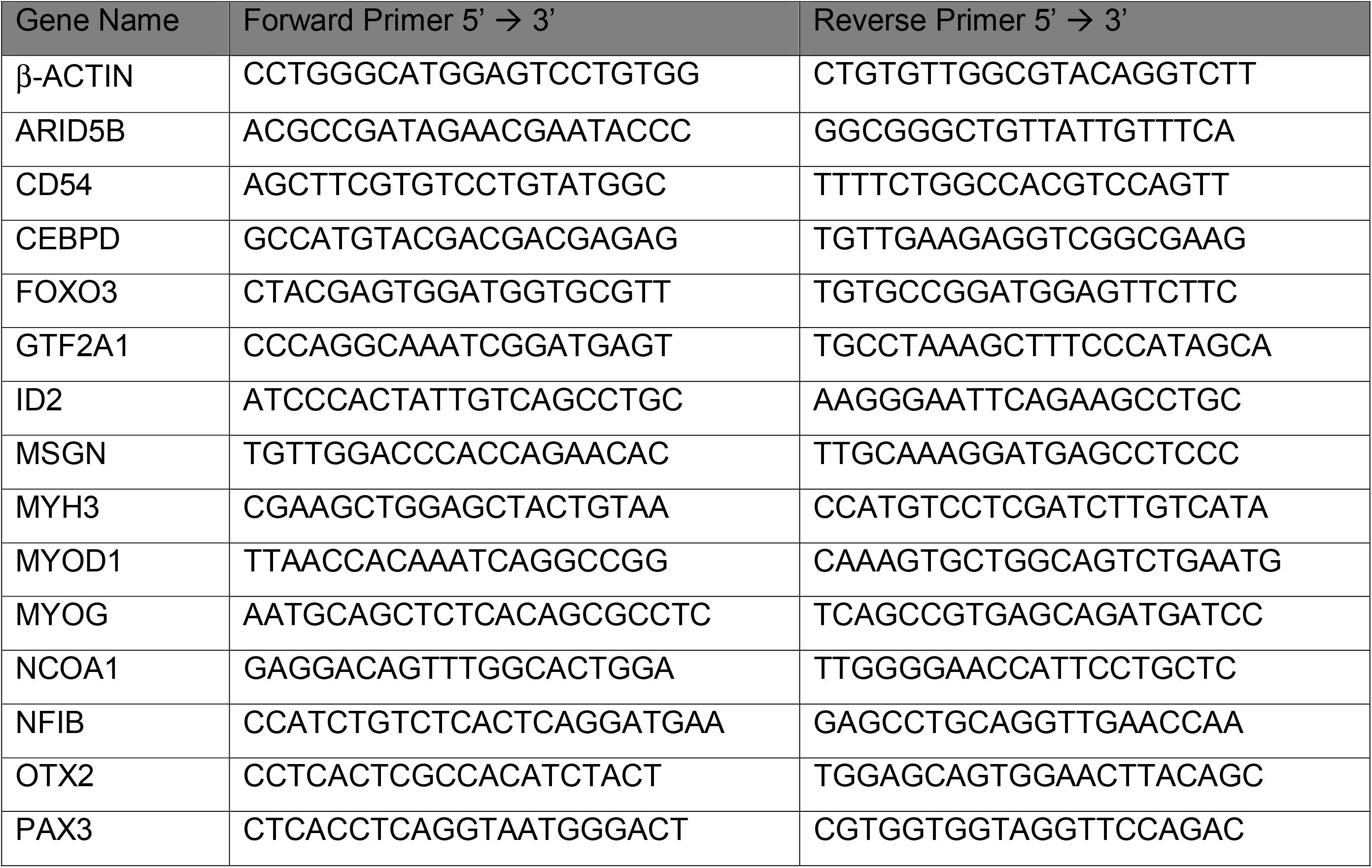
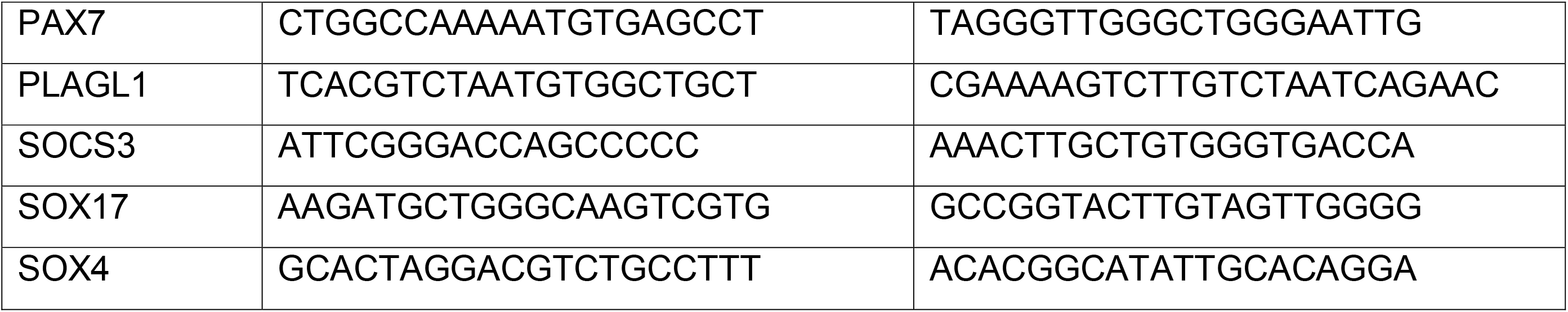

### Immunofluorence staining

Cells grown on BME-coated 24-well tissue culture plates (Greiner Bio-One) were fixed with 4% PFA for 10 minutes at room temperature, and permeabilized in 0.5% Triton X-100 for 10 minutes then blocked in 4% BSA (blocking buffer) for 30 minutes. Incubation with the primary antibodies diluted in blocking buffer was performed overnight. Primary antibodies: mouse anti-Pax7 (PAX7, DSHB, 1:20), mouse anti-myosin heavy chain (MF20, DSHB, 1:20), rabbit anti-Myog (sc-576, Santa Cruz, 1:200), mouse anti-desmin (MA5-13259, Thermo Scientific). Secondary antibodies: AlexaFluor goat anti-mouse 488 (Invitrogen A11001), AlexaFluor goat anti-rabbit 546 (Invitrogen A11010), Goat anti-mouse IgG, Fc subclass 1 specific (Jackson Immuno Cat # NC0469362), Goat anti-mouse IgG, Fc subclass 2b specific (Jackson Immuno Cat # NC0266980). Nuclei were counterstained with Hoechst 33258 pentahydrate (bis-benzimide) 2 µg/ml (Life Technologies Cat #H3569). Cell proliferation was measured by EdU incorporation and Click-iT chemistry (Life Technologies Cat # C10340). Cell death was measured by TUNEL assay (Roche Cat # 11 684 795 910). Images were acquired with fluorescence microscopy utilizing the Leica DMi8. Images were acquired with 10x and 20x objectives. Fields reported in figures are representative of all examined fields.

### Immunofluorescence Quantification

Quantification was perform using FIJI software. Images from each channel were stacked to adjust brightness and contrast. Background threshold was set using the Otsu algorithm and set to 0. Nuclear signal intensity was measured using Analyze Particle function and normalized to nuclear area.

### scRNA-seq Analysis

scRNA-seq dataset were previously described^8^. Digital gene expression (DGE) matrices were downloaded from GEO (GSE147457). Downstream analysis was performed using R (v.4.3.2). Quality control, filtering, data clustering, visualization, and differential expression analysis were performed using Seurat v5 (https://github.com/satijalab/seurat)^78^. Datasets were processed, merged and normalized without integration, as according to instruction from Xi et al^8^. “S.Score” and “G2M.Score” were generated for each cell using “CellCycleScoring”. “Stress” score was generated using the Seurat function “AddModuleScore” utilizing the stress genes dataset provided in the supplementary information of Xi *et* al. The merged dataset was then scaled performing “ScaleData”, with “S.Score”, “G2M.Score”, and “Stress” passed into the vars.to.regress variables.

### Developmental trajectory Analysis

Developmental trajectory analysis was calculated using the non-linear dimensional reduction pipeline of PCA-UMAP^79^. Clusters identified with developmental stages based on the distribution and prevalence of individual *in vivo* samples in each cluster.

### Western Blotting

Total protein extracts were prepared from cultures using lysis buffer (50 mM Tris HCl, 100 mM NaCl, 1 mM EDTA, 1% Triton 100X, pH = 7.5) containing protease and phosphatase inhibitor cocktails (Roche). Cell debris was removed by centrifugation and protein concentration was determined using a standard BSA curve (Pierce, Thermo Scientific Cat #23225). Total protein extracts (30 μg) were loaded onto a NuPAGE 4-12% Bis-Tris gel and electrophoresis was performed in MOPS SDS running buffer (Invitrogen). Proteins were transferred to a PVDF membrane (Bio-Rad) and blocked with 5% bovine serum albumin (BSA) in PBST (PBS with 0.1% Triton 100X). Incubation with primary antibodies was performed overnight at 4 °C. The antibodies used were: rabbit anti–STAT3 (Cell Signaling D3Z2G Cat #12640), rabbit anti-pSTAT3-Y705 (Cell Signaling D3A7 Cat #9145), mouse anti-PAX7 (DSHB Cat #PAX7-c), mouse anti-tubulin (Sigma Cat #T5168), rabbit anti-GAPDH (Cell Signaling 14C10 Cat #2118) and HRP-conjugated secondary antibodies (Santa Cruz). Membranes were visualized with enhanced chemiluminescence (Pierce, Thermo Scientific Cat #32106) and developed on film.

### CD54 MACS isolation

Isolation of CD54+ cells was performed as described in Magli et al^51^. Briefly, at D30 of the differentiation protocol, myogenic culture were isolated into single cells, collected, washed with PBS, and filtered through a 70μm strain to remove cell clumps. Single cells were incubated with human blocking IgG (BD Pharmingen Cat #564219), at 1 µl/10^6^ cells in Blocking buffer (PBS with 10% FCS) for 5 minutes on ice. Staining was performed by adding 1 µl of Anti-Human CD54 (ICAM-1) Biotin (eBioscience, Cat #13-0549-82) for 20 minutes on ice. Magnetic isolation was performed using Streptavidin-conjugated microbeads (Miltenyi Cat #130-048-101) following the manufacturer’s instructions on MS columns (Miltenyi Cat #130-042-201).

## Statistical analysis

Data are presented as mean ± SEM. All statistical tests were performed using GraphPad Prism 10. Investigators were not blinded to group allocation or outcome assessment. No samples were excluded from this study.

## Authors’ contributions

LC and AS conceived the study and wrote the manuscript. LC, CS, MTT, and AS designed and performed the experiments. MSF, AF, RBD, ENO provided patients derived fibroblasts and hiPSCs. AS and PLP provided funding support. All authors participated in the scientific discussion and in the review and editing of the manuscript. The authors read and approved the final manuscript.

