## Supplemental Figures for "Modulation of the JAK2-STAT3 pathway promotes expansion and maturation of human iPSCs-derived myogenic progenitor cells"

[1]Sanford Burnham Prebys Medical Discovery Institute, Development, Aging and Regeneration Program, La Jolla, CA 92037, USA; [2]Graduate School of Biomedical Sciences, Sanford Burnham Prebys Medical Discovery Institute, La Jolla, CA 92037, USA; [3]UOL Medical Genetics, University of Ferrara, Ferrara 44121, Italy; [4]Department of Molecular Biology, Hamon Center for Regenerative Science and Medicine, Senator Paul D. Wellstone Muscular Dystrophy Cooperative Research Center, University of Texas Southwestern Medical Center, 5323 Harry Hines Boulevard, Dallas, TX 75390, USA.

[$]Current Address: Howard Hughes Medical Institute, Robin Chemers Neustein Laboratory of Mammalian Cell Biology and Development, The Rockefeller University, New York, NY 10065, USA.

[#] Lead Author

#### Supplementary Figures Legend

**Supplementary Figure 1: Related to figure 1: AG490 mediated inhibition of JAK/STAT3 does not induce apoptosis**

A. Representative immunofluorescence images of myotubes either from healthy or Duchenne patient-derived myoblasts treated with different JAK inhibitors for MF-20 (red), nuclei are counterstained with Hoechst (blue). Scale bar 100µm
B. Differentiation index of myotubes for the conditions shown in (A) n=3 biological replicates.
C. Representative immunofluorescence images of myoblasts stained with Desmin (green), TUNEL (red) nuclei are counterstained with Hoechst (blue). Scale bar 10µm.
D. Quantification of the percentage of TUNEL positive myoblasts shown in (C). n=3 biological replicates.

ns. $p > 0.05$ ; *. $p \leq 0.05$ ; **. $p \leq 0.01$ ; ***. $p \leq 0.001$ unpaired Student's t test.

* over untreated control; # over lower concentration of drug; $ over DMSO – vehicle control.

**Supplementary Figure 2: Related to figure 4: generation and characterization of hiPSC lines from healthy donor and DMDΔ52 patient.**

A. Schematic representation of the genotyping for DMDΔ52.
B. Representative immunofluorescence images of OCT4 (red) in different hiPSCs clones, nuclei are counterstained with Hoechst (cyan), H9 is shown as positive control. scale bar 100µm.
C. qPCR analysis of markers for germline formation in embryoid bodies assay, *OTX2* (Ectoderm), *SOX17* (Endoderm), *MSGN* (Mesoderm), H9 are shown as positive control (n=3 biological replicates).

ns. $p > 0.05$ ; *. $p \leq 0.05$ ; **. $p \leq 0.01$ ; ***. $p \leq 0.001$ (unpaired Student's t test).

**Supplementary Figure 3: Related to figure 4: AG490 treatment leads to increase myogenesis in both healthy and DMD myogenic cultures**

A. qPCR analysis of *PAX7* expression during the myogenic differentiation protocol in healthy (red) and DMDΔ52 (blue) myogenic cultures.
B. Representative immunofluorescence images of MYOG (magenta) at D30 of differentiation, nuclei are counterstained with Hoechst (gray). Scale bar 50µm.
C. qPCR analysis of *PAX7, MYOD1* and *MYOG* expression at D30 of the differentiation protocol in a second clone of hiPSCs.
D. qPCR analysis of myogenic factors (*PAX7, MYOD1, MYOG*, and *MyHC*) expression at D50 of differentiation in a second clone of hiPSCs.

N=3 biological replicates.

ns. $p > 0.05$ ; *. $p \leq 0.05$ ; **. $p \leq 0.01$ ; ***. $p \leq 0.001$ unpaired Student's t test for A - C - E - G, One-Way ANOVA for D.

**Supplementary Figure 4: Related to figure 4: AG490 treatment leads to increase myogenesis in both healthy and DMD myogenic cultures independently of DMD mutation**

A. Schematic representation of the genotyping for DMD$\Delta$8/9.
B. Representative immunofluorescence images of PAX7 (green) and MYOG (magenta) at D30 of differentiation, nuclei are counterstained with Hoechst (gray). Scale bar 50µm.
C. Violin plot or the quantification of the nuclear mean fluorescence intensity (arbitrary unit) (median $\pm$ second and third quantile) of the PAX7 IF signal staining in healthy (left panel) and DMD$\Delta$8/9 (right panel) myogenic cultures.

N=3 biological replicates.

ns. $p > 0.05$ ; *. $p \leq 0.05$ ; **. $p \leq 0.01$ ; ***. $p \leq 0.001$ unpaired Student's t test for D, One-Way ANOVA for C.

**Supplementary figure 5: Related to Figure 5: AG490 treatment leads to maturation of *in vitro* derived SMPC**

qPCR analysis of *SOX4, GTF2A1* (STAGE 1/2), *PALGL1* (STAGE 4) and *CEBPD, FOXO3* and *ARID5B* (STAGE 5) expression in CD54 MACS-purified populations.

N=3 biological replicates.

ns. $p > 0.05$ ; *. $p \leq 0.05$ ; **. $p \leq 0.01$ ; ***. $p \leq 0.001$ unpaired Student's t test.

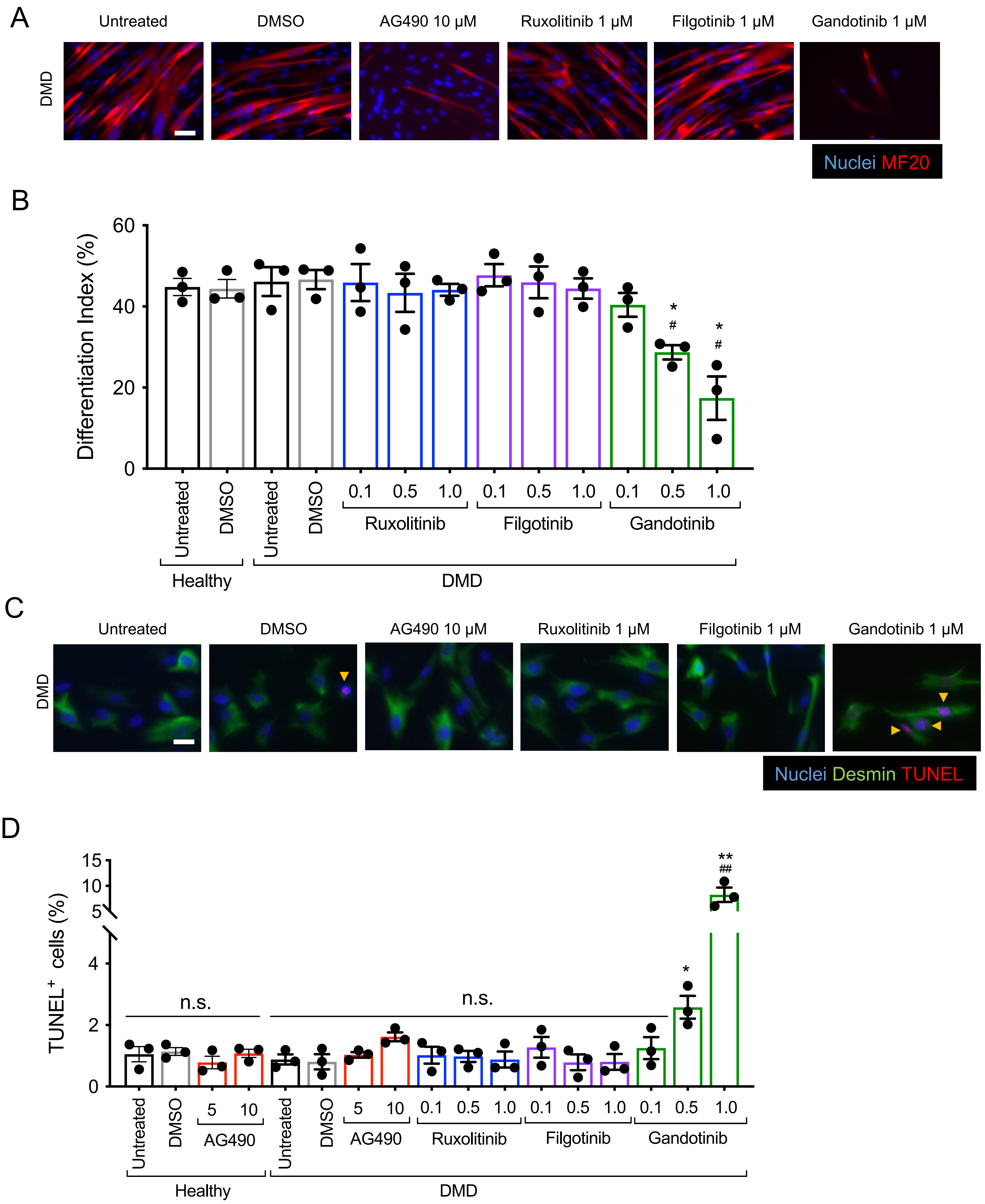


**Supplementary Figure 1**

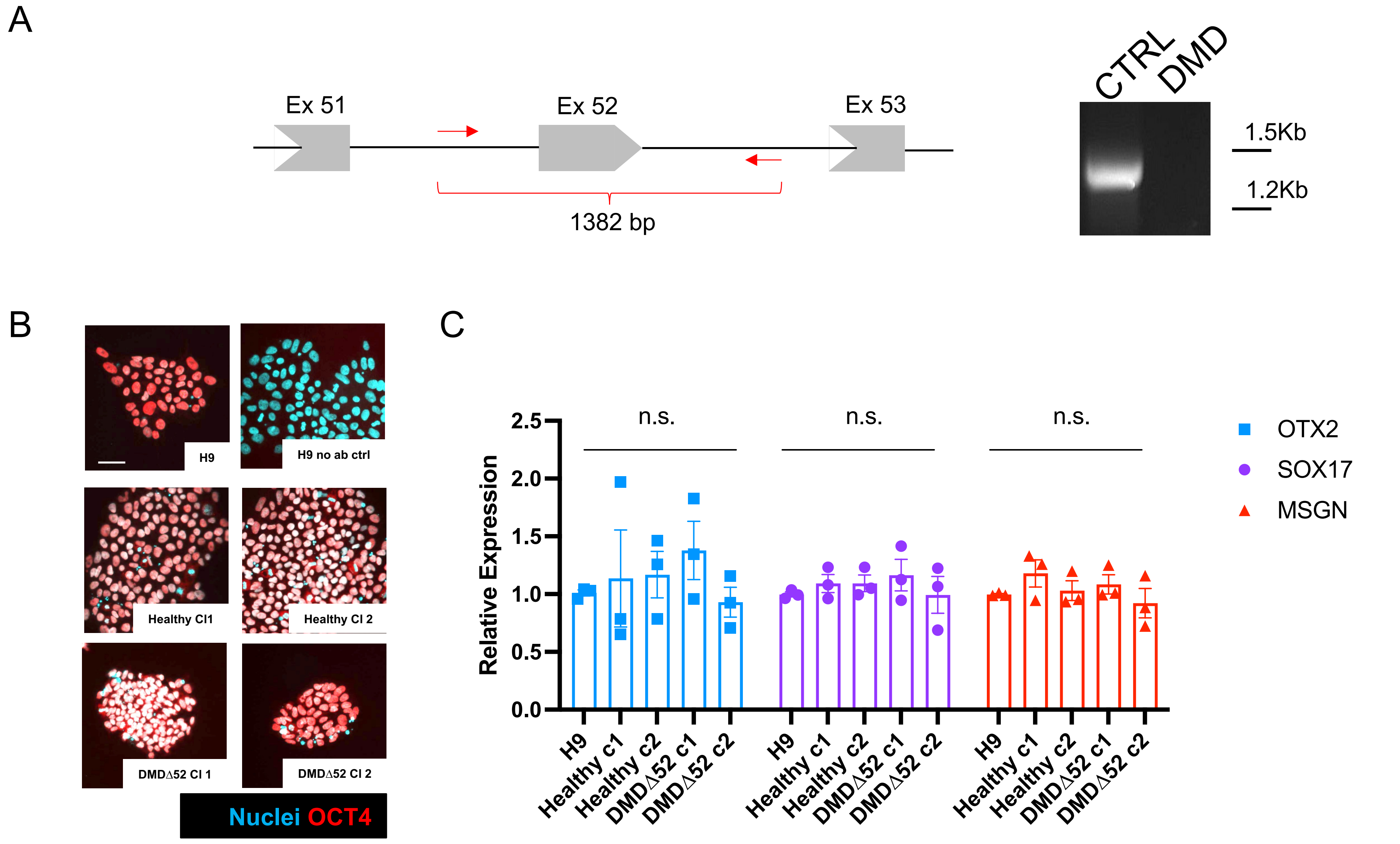


**Supplementary Figure 2**

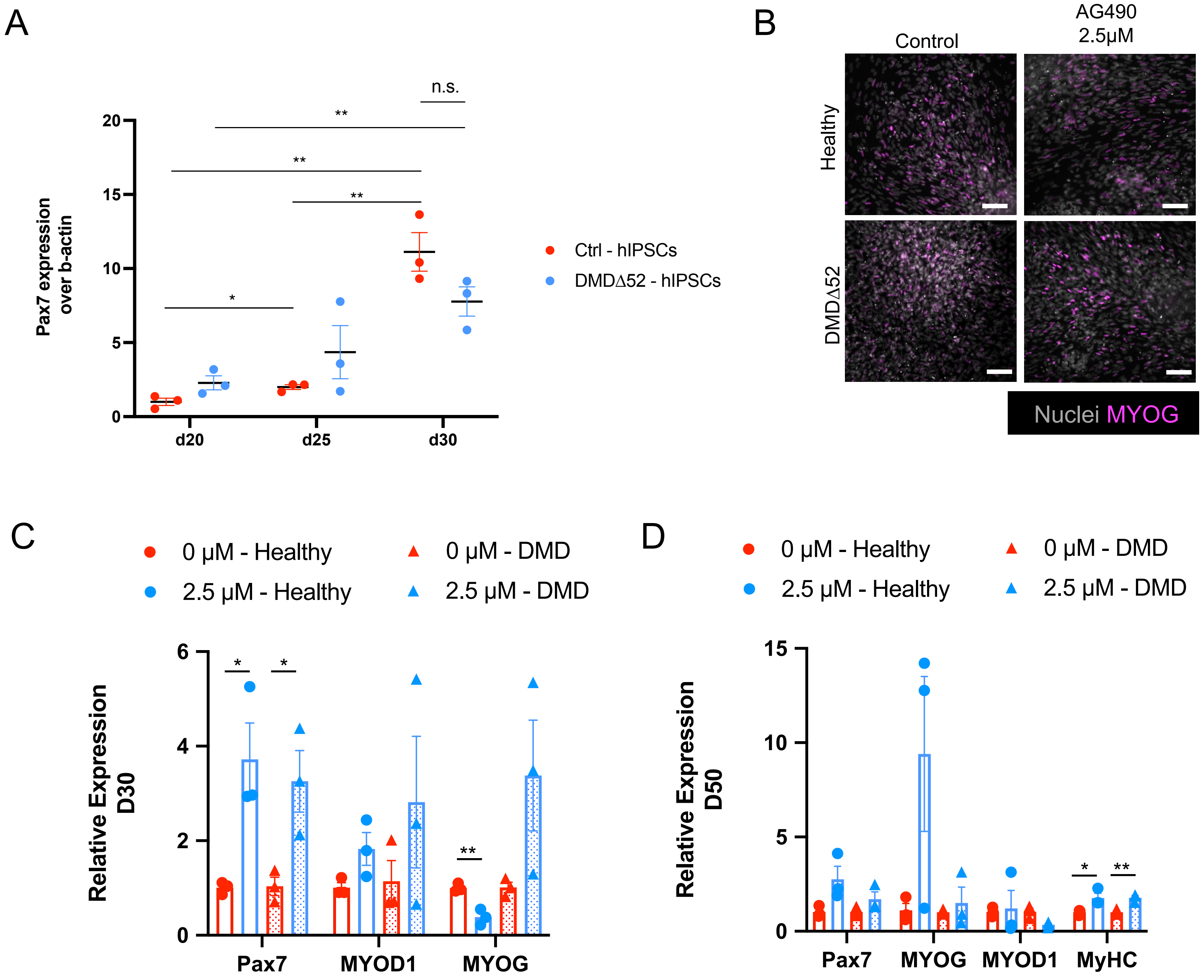


**Supplementary Figure 3**

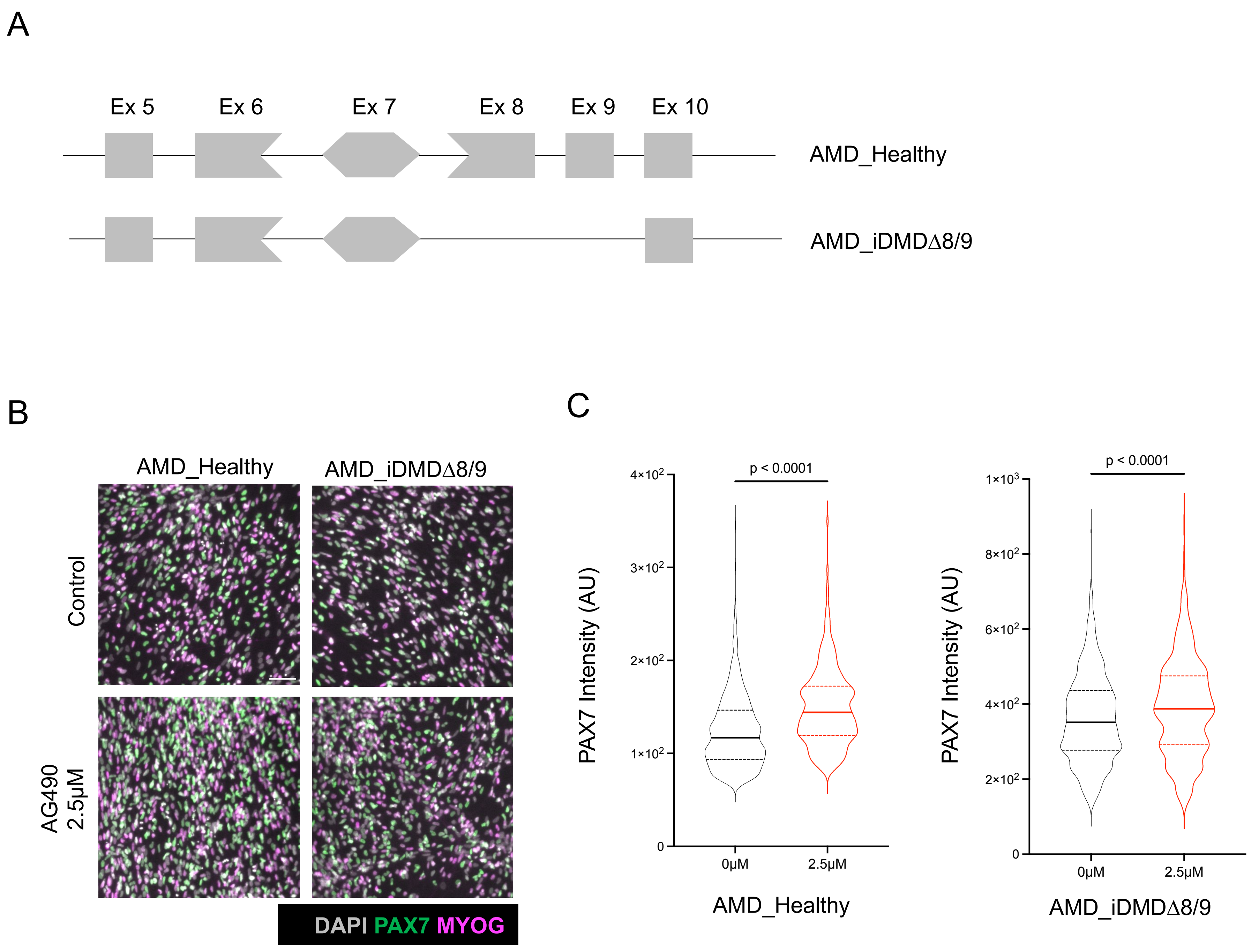


**Supplementary Figure 4**

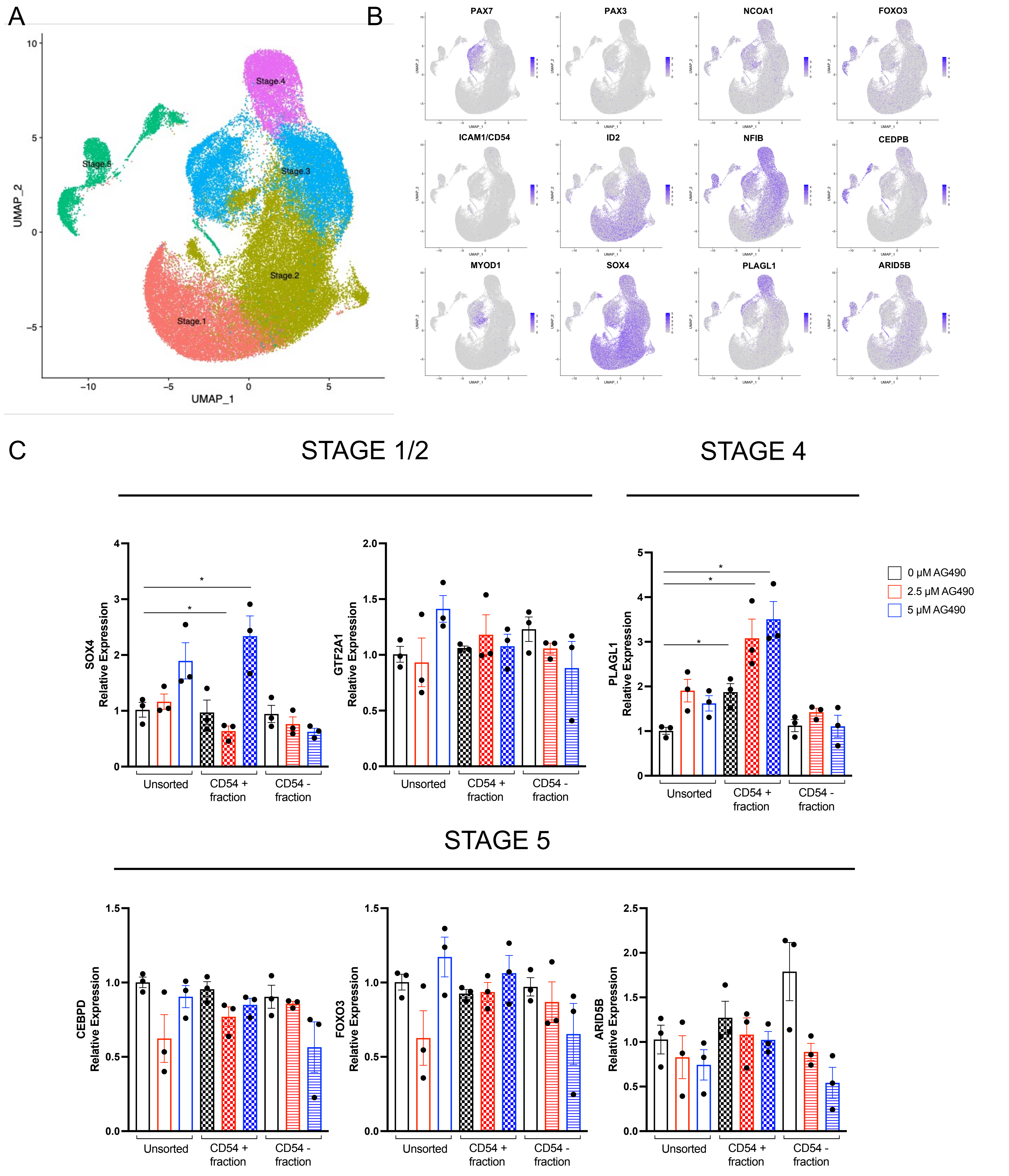


### Supplementary Figure 5
